# Exploring noise in TCR specificity data through supervised learning

**DOI:** 10.64898/2026.10.02.756166

**Authors:** Sebastian Nymann Deleuran, Yogesh Basavaraju, Rūta Matulevičiūtė, Kristoffer Haurum Johansen, Sine Reker Hadrup, Morten Nielsen

**Affiliations:** Department of Health Technology, Technical University of Denmark, DK-2800 Lyngby, Denmark

## Abstract

Accurate prediction of T cell receptor (TCR) specificity can potentially accelerate development of novel immunotherapies, yet public databases used to train these models contain substantial label noise. This uncertainty complicates model evaluation and limits the reliability of predicted TCR-pMHC interactions. In this work, we present a simple supervised machine learning denoising scheme to identify low-quality labels in TCR specificity data. We evaluated the framework using the TCRvdb validation dataset and further corroborated it via structural modeling with AlphaFold 3. Here, our denoising approach successfully identified non-validated TCRs, achieving an average MCC of 0.47 for filtering non-validated examples while retaining high sensitivity for validated examples. Applying our denoising approach to the complete NetTCR training dataset, we observed that data associated with known noisy attributes were significantly enriched in the filtered subset. Furthermore, TCR-pMHC complexes identified as noise consistently produced lower AlphaFold 3 confidence scores, providing independent evidence of misannotation. Experimental validation of 30 selected TCRs further supported the approach, with 80% of TCRs predicted as noise failing to demonstrate functional reactivity. Our findings demonstrate that current performance metrics may be obscured by dataset inaccuracies and provide a method for refining training data in the absence of gold-standard benchmarks.

## Introduction

The interaction between the T cell receptor (TCR) and the peptide-major histocompatibility complex (pMHC) represents one of the cornerstones of adaptive immunity [1]. As the experimental discovery of TCR specificity remains cumbersome, in silico approaches for predicting TCR-pMHC binding represent a valuable asset for the development of T cell immunotherapies [2], [3]. A wide range of methods have been developed to solve this task, ranging from simple sequence similarity-based approaches to large deep learning models leveraging structural information [4], [5], [6], [7], [8], [9], [10]. Such approaches typically draw upon TCR specificity data derived from public repositories such as IEDB, VDJdb, and McPas-TCR, with many incorporating various degrees of unsupervised pretraining steps that leverage unlabeled bulk TCR sequencing data [11], [12], [13], [14], [15], [16], [17], [18]. Agnostic to the training paradigm or modeling approach, all these models must be evaluated on annotated TCR-pMHC binding data, which underlines how essential and intertwined public databases are to the development of TCR-pMHC binding prediction methods. However, recent work by Messemaker et al. has demonstrated, for a subset of VDJdb, that a high proportion of TCR sequences cannot be validated with respect to their annotated pMHC specificities in subsequent binding assays [19]. While it has already been known that databases of TCR-pMHC interactions contain significant proportions of incorrectly annotated specificities, the high proportion of noise demonstrated by Messemaker et al. was unexpected [20]. This brings into question the conditions under which we train and evaluate models for predicting TCR-pMHC binding when neither training nor testing data can be trusted to provide accurate estimations of generalization errors. While methods for predicting noise in high-throughput assay data have already been proposed, post-hoc data denoising on public domain data has, to our knowledge, hitherto not been applied and evaluated extensively [21], [22], [23]. In this work, we propose a simple label noise prediction scheme for identifying possible low-quality positive labels in TCR specificity data, building on work from Jensen and Nielsen [5]. We demonstrate that NetTCR, a model trained on data from the public domain, is capable of identifying noisily labeled data in the Messemaker et al. dataset, and how filtering these examples from the NetTCR dataset changes predictive performance on test data contained in this subset. Additionally, we apply the denoising scheme to the full training dataset of NetTCR, and showcase that filtered data tend to generate low-confidence structural models when modeled with AlphaFold 3 [24].

## Methods

### NetTCR training dataset

The dataset used for training NetTCR was composed of data retrieved from IEDB, VDJdb, and a dataset from 10x Genomics denoised with the ITRAP method as described in Povlsen et al. [21]. The data were processed in accordance with the pipeline described in Jensen and Nielsen [5]. Briefly, quality filtering was performed to ensure complete CDR3 sequences and V/J gene annotations for each observation.

Full-length α-and β-chain sequences were reconstructed from their CDR3 sequence and V/J gene annotations using Stitchr [25]. CDR1, CDR2, and CDR3 sequences were identified with ANARCI using the IMGT scheme [25], [26].

Redundancy reduction was performed in a per-peptide manner using a Hobohm-1-based approach [27]. Here, for each peptide, a kernel similarity method was used to compute similarity between TCRs based on their average CDR3α and CDR3β sequence similarity [28]. If a TCR had more than 0.95 kernel similarity to an already observed TCR, the data point would be removed. The same approach was used to identify and group similar TCRs during dataset partitioning. Finally, negative examples were generated by deliberately mismatching TCRs and pHLA complexes. For each positive data point, five negative samples were created by pairing the pHLA with TCRs drawn from other data points within the same data partition. To reduce the likelihood of sampling cross-reactive binders, TCRs were only sampled from data points whose associated peptide differed from the original peptide by a Levenshtein distance greater than three. For information on the training data peptide distribution, we refer to Supplementary Figure 1.

**Figure 1:**
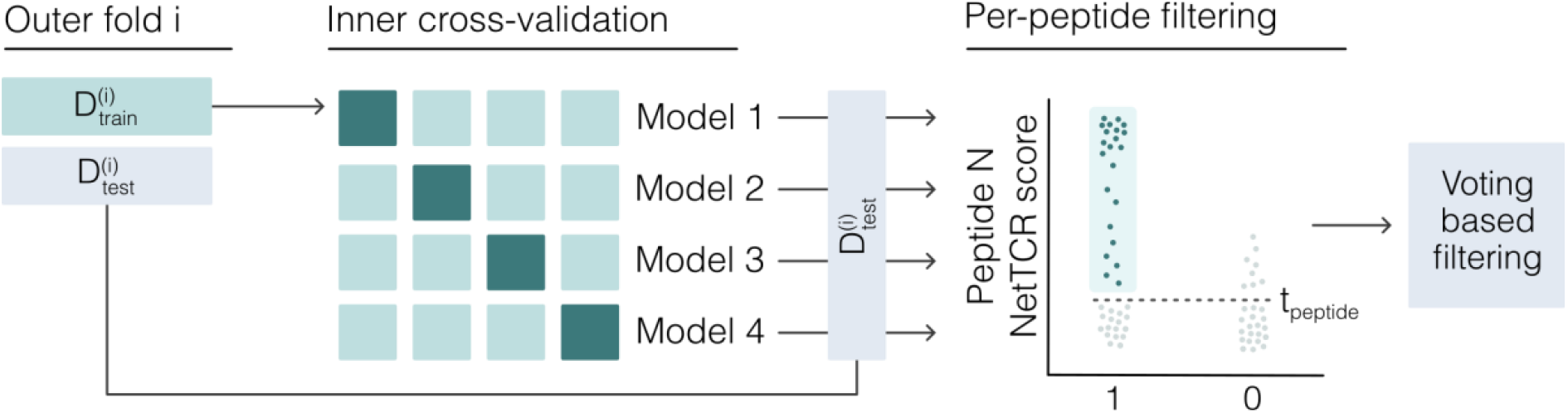
Cross-validation-based data filtering scheme. Our data filtering approach is based on a nested cross-validation framework. For each outer fold *i*, four models are trained on the associated inner folds. These models generate predictions for the outer fold test set. For each peptide, we determine a prediction score threshold defined as the Nth percentile of scores among its negative data points, and use this threshold to filter positive data points. Each model votes to keep or filter data points based on this criterion. The resulting per-peptide filtered subsets are concatenated, and negative data is resampled within each data partition.

### TCRvdb dataset

The dataset used to quantify label noise prediction performance was constructed from the TCRvdb dataset, using the 01-05-2025 version [19]. We followed the authors’ method of generating binary labels for validated and non-validated data points by labeling TCRs associated with a significant (*p*<1e-5) log2 fold change in CD69 expression as validated and those associated with non-significant changes as non-validated. This resulted in 132 and 54 validated and non-validated data points for the peptide GLCTLVAML and 267 and 172 validated and non-validated data points for the peptide YLQPRTFLL.

When quantifying the concordance between data filtered using the various denoising approaches and TCRvdb labels, we only considered the overlap between data in the NetTCR training dataset and TCRvdb. This amounted to 86 and 35 validated and non-validated examples for GLCTLVAML and 257 and 130 validated and non-validated examples for YLQPRTFLL. For the remaining analyses, we used the complete TCRvdb dataset.

### Pan-specific NetTCR models

The denoising framework was showcased using a modified version of previous NetTCR models, implemented in PyTorch [5], [29], [30]. NetTCR is a convolutional neural network consisting of seven one-dimensional convolutional blocks, each operating on a BLOSUM50-encoded segment corresponding to the CDR1α, CDR2α, CDR3α, CDR1β, CDR2β, CDR3β, and peptide, respectively. Each block comprises five one-dimensional convolutional layers with kernel sizes of 1, 3, 5, 7, and 9, respectively, each using 32 filters followed by a max-pooling operation. Each block, therefore, produces five scalar outputs. The outputs of all convolutional blocks are concatenated, passed through a dropout layer with a dropout rate of 0.6, and then passed through a three-layer fully connected network to produce a single scalar representing the predicted binding score. The model is trained in a supervised classification setting using the binary cross-entropy loss function. Optimization is performed using the Adam optimizer with a learning rate of 5e-4, without the use of a learning rate scheduler or L2 regularization. Models were trained for 200 epochs without the use of early stopping. Additionally, to emphasize underrepresented peptides, we employed a sample-weighting scheme in which the loss for each example was scaled inversely to the frequency of its peptide in the dataset, following the approach described by Jensen and Nielsen [5]. The frequency was computed within dataset partitions.

NetTCR models were trained and evaluated using a nested 5-fold cross-validation setup. The dataset was first divided into five partitions (we refer to the section on the NetTCR training dataset). We then performed nested cross-validation with an outer 5-fold loop and an inner 4-fold loop. For a given outer fold *i ∈* {1,…,5}, one partition was held out as the test set, denoted 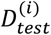, while the remaining four partitions formed the outer training set, denoted 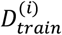.

Within each outer training set, an inner 4-fold cross-validation was performed. For a given inner fold *j ∈* {1,…,5} \ {i}, one partition of 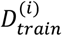 was used as the validation set, denoted *D*^(*ij*)^, and the remaining three partitions formed the inner training set, denoted 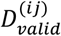. To obtain prediction scores for 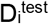, the four models trained during the inner cross-validation were applied to 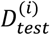, and the final test set predictions were obtained by averaging the outputs of these models. When applied in our denoising setup, the scores were not averaged and were instead used in an ensemble voting scheme.

### Data denoising scheme

In order to identify data points with noisy labels, we employed a denoising scheme based on our nested cross-validation setup (Figure 1).

For an outer fold *i*, we held out the corresponding test partition 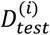 and trained *k* - 1 models, with each possible training and validation data partitioning. For *k* = 5, this yielded four models trained on distinct data subsets. In a per-peptide manner, we considered the set of negative examples in 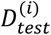, denoted 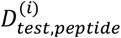, and identified the score threshold corresponding to the Nth percentile of this negative score distribution, denoted *t*_*peptide*_. Under the assumption that positive examples scoring below this threshold exhibit behavior inconsistent with what the model learned during training, we filtered all positive examples in 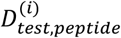 whose cross-validated scores fell below *t*_*peptide*_. Repeating this for all four models, we obtained four votes on whether a data point should be considered noise.

When evaluating the effect of denoising on predictive performance, we held out the test partition from the denoising scheme entirely and instead denoised the inner fold validation partitions. Consequently, only one prediction per data point was available, and we therefore did not employ a voting scheme to identify noisy examples for that analysis. Negative examples were resampled within partitions using the same protocol as was used for the full dataset, in order to avoid having negative data points with TCRs that were sampled from filtered positive data points. NetTCR models were subsequently trained and evaluated on the five datasets resulting from applying this scheme for all *i*, using the nested cross-validation scheme described in the section on *Pan-specific NetTCR models*.

### Enrichment analysis

To analyse the enrichment of features in data filtered using our label noise prediction approach, we computed the log2 fold change in the frequency of each feature in the filtered data, denoted 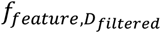, relative to the frequency of those features in the complete data, denoted 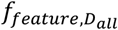, expressed by:

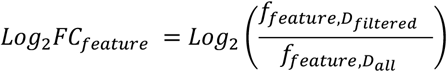

We then computed the statistical significance of each feature’s log2 fold change using Fisher’s exact test for cases where feature counts were low and the Chi-squared test for remaining cases. Low-count cases were defined as those in which any cell of the contingency table had a count below 5. To account for multiple hypothesis testing, we applied the Bonferroni correction to the resulting p-values and used the adjusted p-values to determine significance.

### Structural modeling of TCR-pMHC complexes

TCRα and TCRβ sequences were trimmed to the variable domain using ANARCI, and the MHC sequence was trimmed to the α_1_ and α_2_ domains using an MHC-specific hidden Markov model sequence profile as described in Yin et al. [31]. Structural models were generated using AlphaFold 3. The source code and parameters were obtained from their official sources. In order to reduce the computational cost of generating multiple sequence alignment features, we utilized reduced sequence databases as described in Yin et al. Structural models were created with one seed and ten diffusion samples, yielding ten models per TCR-pMHC complex. Models were ranked on their confidence score defined by confidence = 0.8 · ipTM + 0.2 · pTM + 0.5 · disorder - 100 · has_clash, where ipTM describes the predicted interchain interface accuracy, pTM measures the predicted overall structural confidence, disorder describes the proportion of disorder across the protein, and has_clash is a penalizing term for clashing atoms. The max per-complex confidence scores were used for subsequent analyses.

### Functional screening of selected TCRs

T cell receptors for validation were ordered as gene fragments containing TRBV-TRBC-P2A-TRAV (Twist Bioscience) with BsmBI overhangs and cloned into a lentiviral plasmid containing the TRAC locus expressed under an EF1alpha promoter. Plasmids were transfected into HEK293T cells pre-seeded 24hours prior in 96-well plates using Lipofectamine 3000 (Thermofisher Scientific L3000015). After 48 hours, the lentivirus was harvested by centrifuging the plate at 1500 g × 20 min at room temperature. Then, TCR KO, CD3 KO, CD8+ Jurkat E6.1 were transduced with the virus in 96-well plates. Following 7 days of culture, the cells were stained with pMHC tetramers (generated by exchanging the peptide-of-interest to the respective MHC complex and tetramerizing them using APC-and BV421-conjugated streptavidin complexes) for detecting the cognate TCR at RT for 30 min, followed by antibody mix containing anti-CD3 FITC (BD Biosciences 555332), anti-CD8 BV480 (BD Biosciences 566121) and a dead cell marker (LIVE/DEAD fixable Near-IR; Invitrogen L10119) at 4°C, 30 min in FACS buffer (PBS+2%FBS). Following 2x wash in FACS buffer, cells were acquired on a Fortessa 2 flow cytometer (BD Biosciences). Data were analysed for pMHC tetramer binding within the CD3+ population using FlowJo (V10.0.0).

## Results

In this section, we first evaluate the ability of the NetTCR-based supervised denoising framework to identify potentially mislabeled TCR-pMHC interactions using the independently validated TCRvdb dataset. We then characterize the properties of the filtered data through enrichment analyses and structural modeling with AlphaFold 3. Next, we assess the impact of denoising on NetTCR predictive performance. Finally, we provide experimental validation of selected TCRs predicted to represent noise.

### Learning biologically relevant features in the context of label noise

In order to identify data points that were likely to be incorrectly labeled, we implemented a supervised cross-validation-based label noise prediction scheme. In short, for each peptide, we trained a NetTCR model using nested cross-validation, and then computed a score threshold corresponding to the Nth percentile of scores for that peptide’s negative data points. All positive examples that were scored lower than that threshold were filtered and labeled as noise (for details on the data, data partitioning, models, and model training, we refer to the Methods section). We considered the 75th, 80th, 85th, 90th, 95th and 99th percentile score thresholds. For each data point, we obtained four scores as a result of the nested cross-validation scheme. We observed that some data points were consistently predicted to have low scores across models, indicating agreement that these examples resemble negatives. In contrast, high-variance examples were likely to represent ambiguous interactions rather than mislabeled data. Therefore, we use the four scores as an ensemble voting system, filtering data points with increasing degrees of agreement between models.

To quantify the performance of the label noise prediction scheme, we used data from the TCRvdb dataset containing TCR specificity data for the two peptides YLQPRTFLL (YLQ) and GLCTLVAML (GLC). Briefly, this data specifically describes whether data points obtained from the VDJdb could be validated in a secondary set of T cell activation assays, which provides a ground truth for evaluating denoising performance [19]. We quantified the denoising performance using a Matthews correlation coefficient describing the concordance between data points filtered with our label noise prediction scheme and data points that were not validated in the T cell activation assays.

Visualizing the distribution of scores for positive data points for YLQ and GLC reveals a correlation between score and validation status in the TCRvdb dataset (Figure 2). Particularly for YLQ, we see a clear bimodal distribution of scores, where data that is scored highly corresponds to validated data points in TCRvdb, and the data in the low-scoring mode are abundantly non-validated. For the latter group, scores fall within the range of the scores of the negative data points for that particular peptide, providing a clear rationale for our denoising approach. The separation is less clear for GLC, though validated examples tend to be scored higher than non-validated examples. We note that the clearer separation of the YLQ data may be due to the uniquely high sequence similarity of some of the validated TCRs (Supplementary Figure 2). To evaluate the performance of the noise prediction framework, we can frame the task as a classification problem, where we quantify how well the noise prediction scheme aligns with the validated and non-validated labels. Applying the filtering scheme on the TCRvdb subset and increasing the filtering score threshold improves filtering MCC for both peptides, with the strictest filtering threshold computed from the 99th percentile of negative scores and no consensus voting, yielding an MCC of 0.6 and 0.18 and an F1 of 0.73 and 0.55 for YLQ and GLC, respectively (Supplementary Figure 3). While this filtering configuration for these particular peptides yields the highest MCC scores and captures the most noise, this threshold results in a very large number of TCRs being discarded, retaining only 71% and 42% of validated examples. Therefore, for the sake of maintaining high sensitivity and not filtering substantial amounts of clean data, we find that using the 90th percentile negative score as the filtering threshold is a good compromise. Additionally, while using ensemble voting reduces the proportion of noise detected, particularly for GLC, it improves sensitivity from 0.71 and 0.42 to 0.91 and 0.66 for YLQ and GLC, respectively.

**Figure 2:**
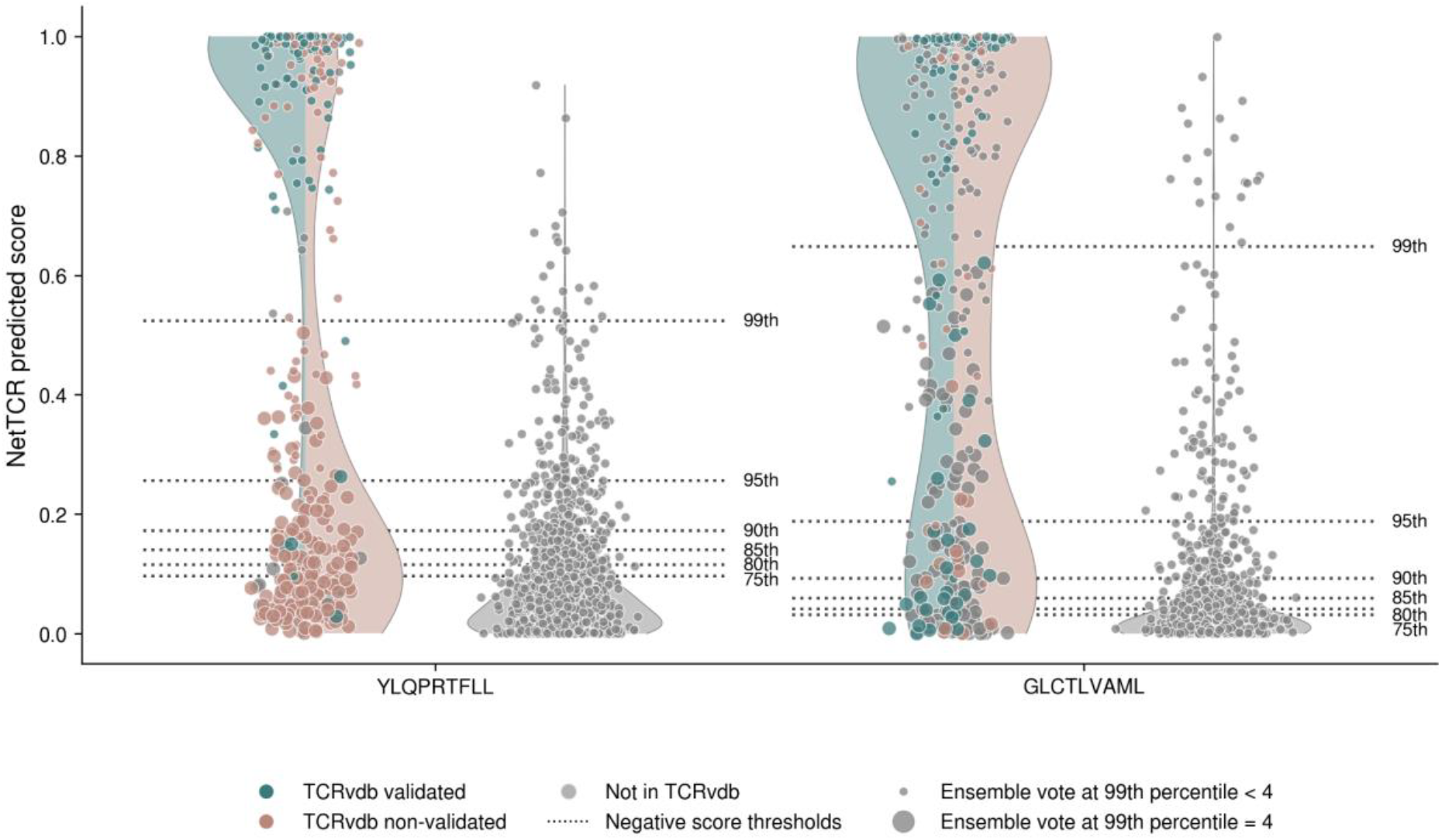
Distribution of NetTCR scores for validated and non-validated TCRs. NetTCR score distribution for positive and negative data points for peptides YLQ and GLC. Data points are colored according to whether or not they are found in the TCRvdb dataset and their validation label. Negative score threshold values for the different percentiles are visualized with dotted lines. Dot size indicates ensemble voting agreement when filtering at the 99th percentile.

**Figure 3:**
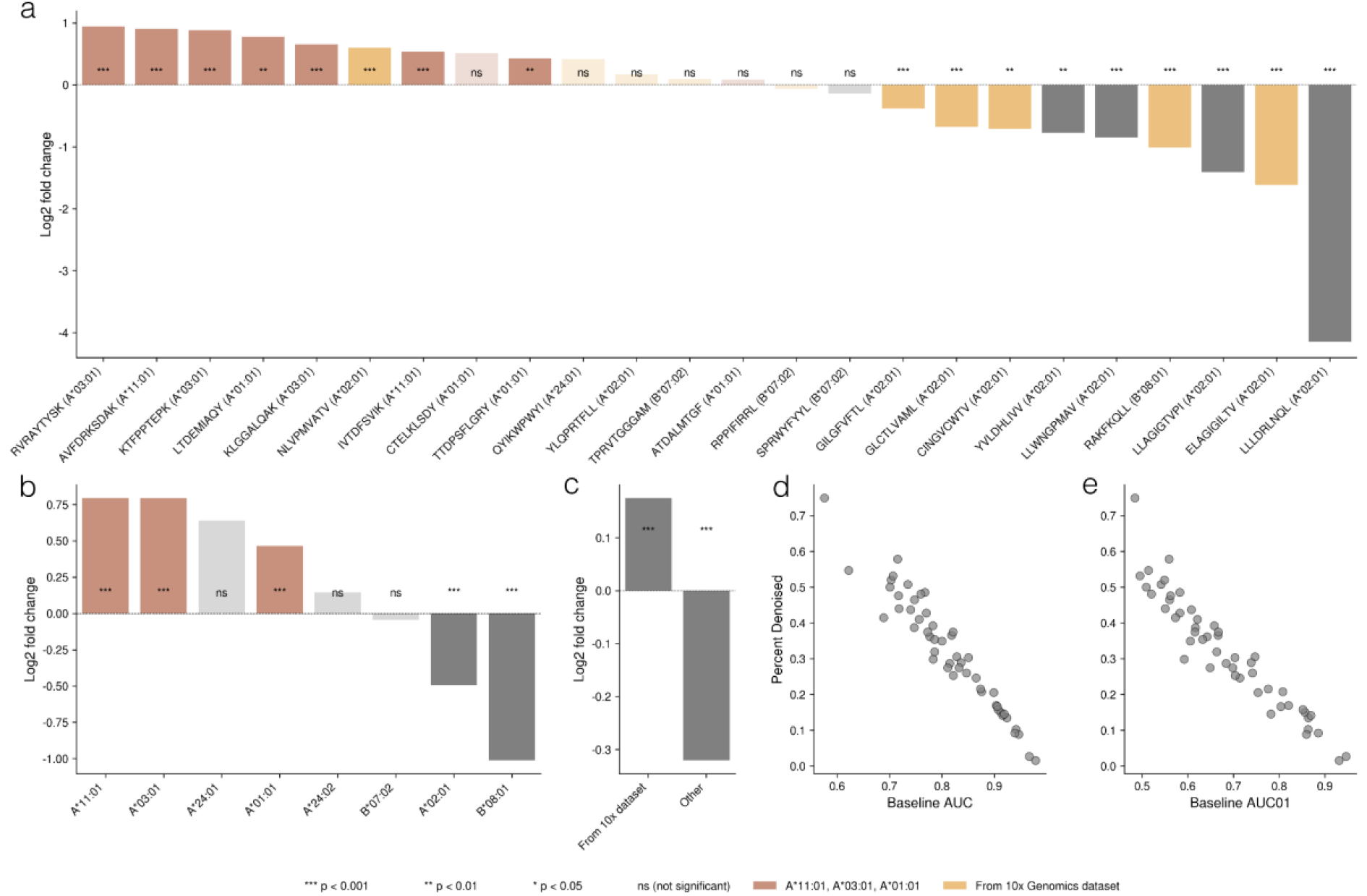
Enrichment of various attributes in filtered data. Data were filtered using a score threshold corresponding to the 90th percentile of negative scores for each peptide. Only data for peptides with more than 100 positive data points were included in the analysis. For peptides, HLA allele features, log2 fold changes of frequencies in the filtered data versus frequencies in the complete data were computed. (a) Log2 fold changes of peptide frequencies in the filtered versus non-filtered data. Red indicates that a peptide is restricted to HLAs known to be noisy in high-throughput assays, and orange indicates that data for a given peptide in part originates from 10x Genomics-derived datasets [23]. (b) Log2 fold changes of HLA allele frequencies in the filtered versus complete data. Red indicates that the allele is known to be noisy in high-throughput assays [22]. (c) Log2 fold changes of data source frequencies. (d) and (e) Per-peptide AUC and AUC01 versus the percent of data denoised using the 90th percentile negative score threshold and full ensemble voting agreement.

### Known sources of noisy data are overrepresented in filtered data

To further evaluate the soundness of our approach, we examined the properties of the data identified as noise in the NetTCR dataset, using the 90th percentile negative score threshold and full ensemble voting agreement. As expected, due to the nature of the noise prediction scheme, peptides with low NetTCR predictive performance are heavily filtered, filtering as much as 40-60% of positive data points for peptides with AUCs below 0.75 (Figure 3d and 3e).

We then examined the enrichment of peptide and HLA allele features in the filtered data by computing the log2 fold change of the frequencies of these features in the filtered data and the complete data (Figure 3a, 3b, and 3c). We opted to only consider data for peptides with more than 100 positive data points in order to improve the statistical power of the analysis. Interestingly, peptides that are enriched in the filtered data coincide with peptides presented by MHC molecules that are known to be associated with high degrees of noise [22], [23]. Specifically, we see an enrichment of A*01:01, A*03:01, and A*11:01 restricted peptides and peptides where TCRs are derived mainly from 10x Genomics datasets, i.e., peptides such as RVR, AVF, KTF, and LTD, with log2 fold changes of 0.94, 0.91, 0.88, and 0.78. This concordance suggests that the denoising scheme likely captures actual noise in the dataset and that biologically irrelevant data points are being filtered from the data.

To further examine the properties of the denoised data through an orthogonal method, we created structural models of TCR-pMHC complexes for peptides that were either over-or underrepresented in the filtered data. For each complex, we generated ten structural models using AlphaFold 3 and used the highest per-complex confidence scores for the analysis (for details, refer to Methods) (Figure 4a, Supplementary Figure 4, Supplementary Figure 5). Here, we observe that for the three peptides (RVRAYTYSK, AVFDRKSDAK, and KTFPPTEPK) with the highest degree of filtering, confidence scores generally fall below 0.7, indicating poor model quality. For less filtered peptides, the disparity between confidence scores for filtered and non-filtered examples grows (Figure 4b). With the intuition that true TCR-pMHC pairs will generally yield higher confidence structural models, these results thus suggest that the filtered data, to a high degree, share a lower likelihood of being binders compared to the non-filtered data [9], [19], [32].

**Figure 4:**
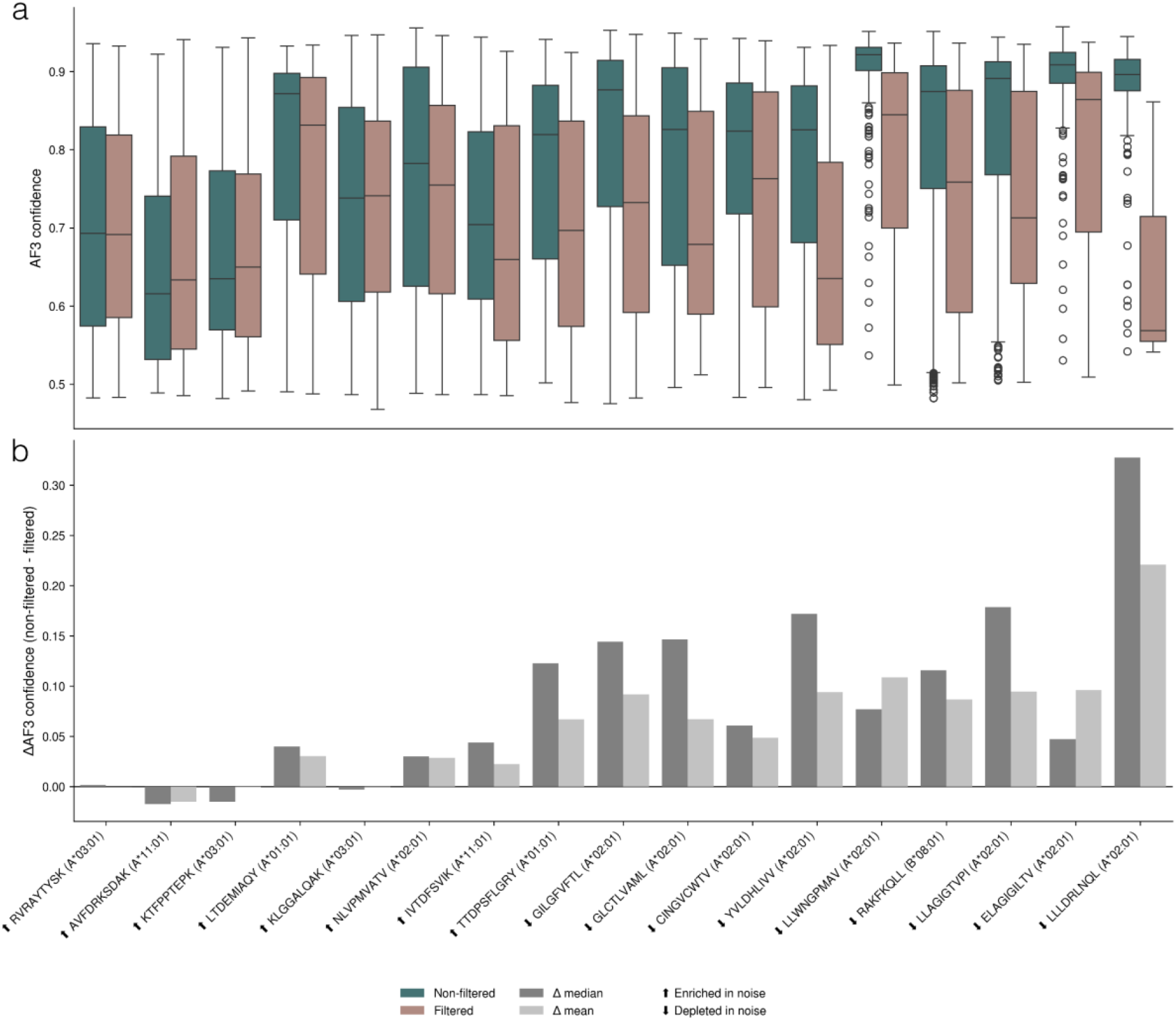
Distribution of AlphaFold 3 confidence scores for enriched and depleted peptides. All positive data points for peptides that were significantly enriched or depleted in denoised data were modeled with AlphaFold 3, and their confidence scores were max-pooled. Peptides are sorted by their log2 fold change of frequencies in the filtered data versus frequencies in the complete data. (a) Confidence scores for filtered and non-filtered data points. This analysis shows that TCR-pMHC complexes for peptides overrepresented in filtered data tend to be modeled with lower confidence. (b) The difference in mean and median between filtered and non-filtered data points.

**Figure 5:**
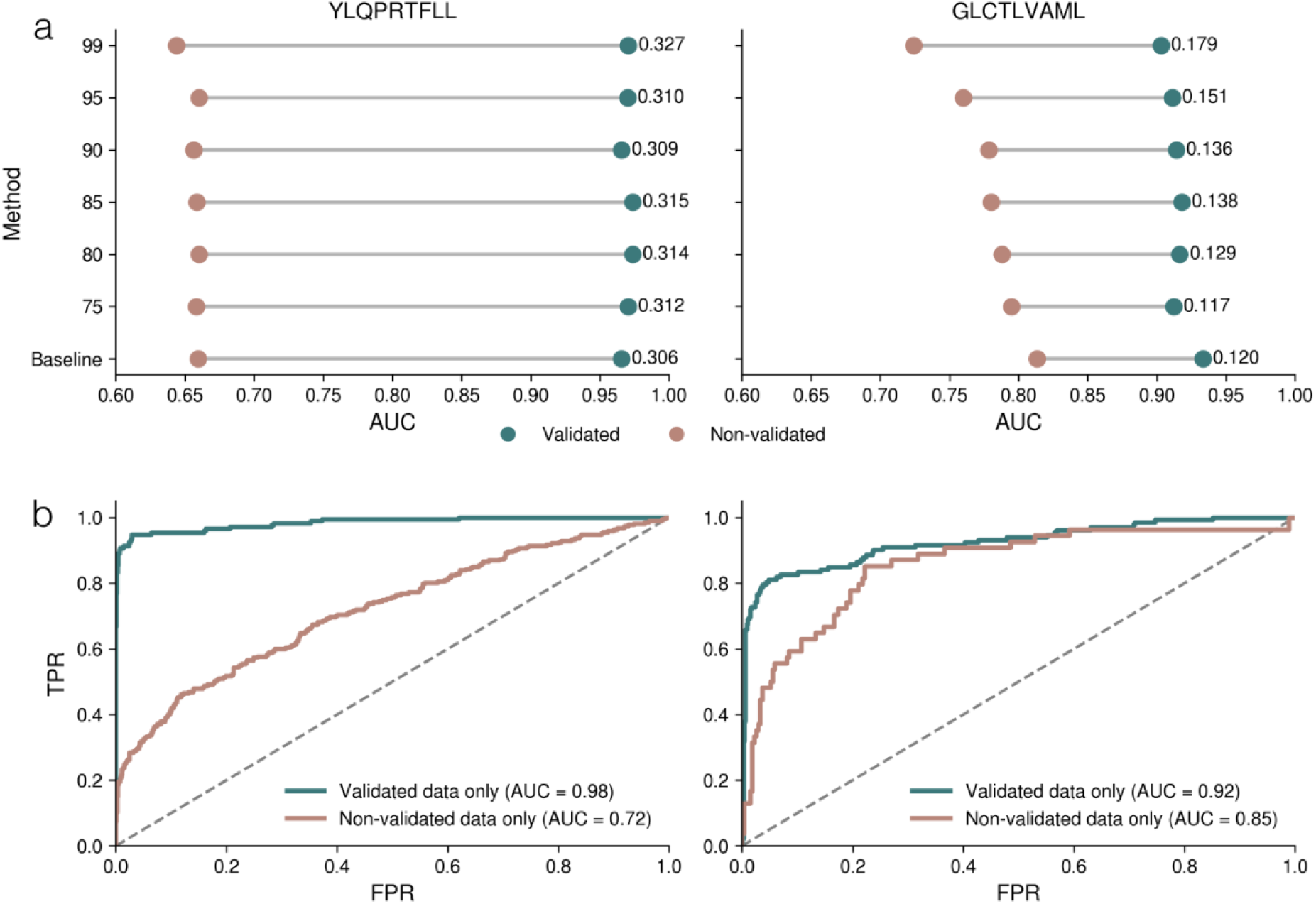
The effect of re-training NetTCR on denoised training data. (a) Predictive performance on validated TCRs versus mismatched negatives and non-validated TCRs versus mismatched negatives, trained on datasets denoised with a range of negative score thresholds. Numbers indicate ΔAUC between the two AUC values. (b) Predictive performance for NetTCR models, trained exclusively on either validated or non-validated TCRs and mismatched negatives. Models were trained on data for a single peptide at a times.

### False positive data with learnable signal challenges NetTCR

We then moved on to investigate the behavior of a NetTCR model when trained on a denoised dataset. Here, we show the performance of NetTCR trained on datasets constructed at different denoising thresholds for GLC and YLG and report the performance as the AUC for the validated and non-validated TCRs versus mismatched negatives (Figure 5a, Supplementary Figure 6). For the subset of validated TCRs, the figure shows nearly constant performance for the models trained on the different denoised datasets. This observation suggests that the NetTCR method, to a very high degree, is robust to noise and is capable of capturing the shared signature in TCR binding data even in the presence of such noise. For GLC, a small performance drop in the validated TCRs is observed as the filtering threshold increases. This reflects the fact that the denoising for this peptide is less accurate compared to YLQ and hence results in a certain proportion of validated binders being removed, resulting in a reduction in the learnable training data signal. For non-validated TCRs, the performance for the YLQ data remains close to constant at the different levels of denoising, whereas the performance for the GLC data displays a consistent performance drop for the models trained on the denoised data. This finding points towards the non-validated GLC TCRs sharing a learnable signal that is captured in the baseline model and is gradually removed as part of the data denoising. This, in contrast, is not the case for the non-validated YLQ TCRs. These conclusions are further supported by performance values of a NetTCR method trained and evaluated using cross-validation on either validated or non-validated TCRs for the two peptides (Figure 5b). When training on non-validated TCRs, the AUC for YLQ is substantially lower than that of GLC, suggesting a lesser degree of a learnable signal in the non-validated data. These results highlight the challenges of benchmarking the performance of methods for data denoising and underline the need for “gold standard” data sets, such as TCRvdb.

**Figure 6:**
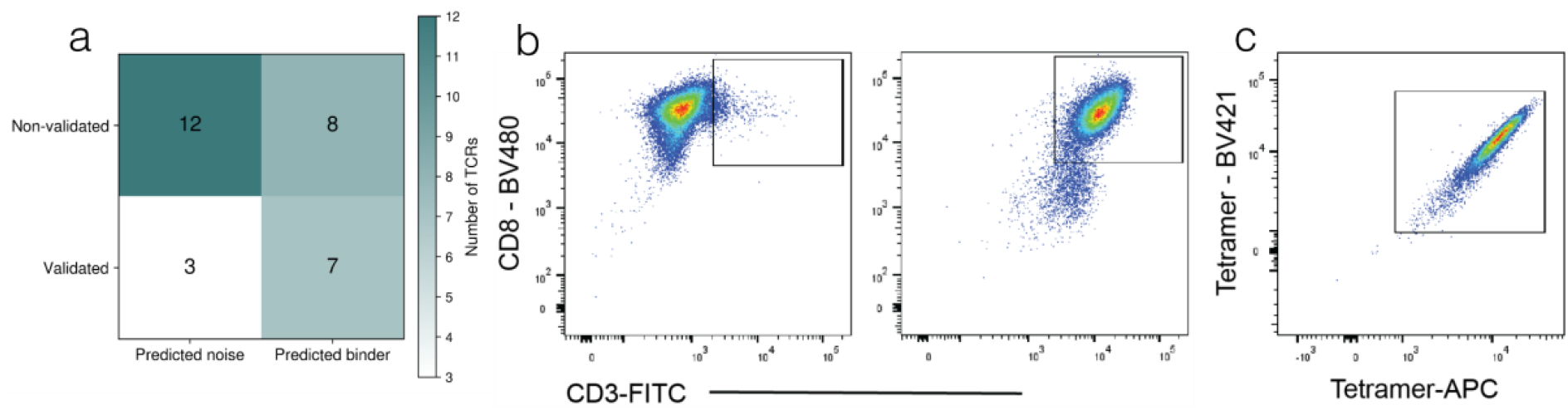
Experimental validation of TCRs predicted to be noise.30. TCRs from public databases for peptides TTDPSFLGRY, YVLDHLIVV, and IVTDFSVIK were scored by NetTCR to be either binders or non-binders. TCRs were transduced into TCR KO CD8+ Jurkat cells using TCR-encoding lentivirus and stained for their cognate pMHC antigen, and their binding status was compared to their predicted labels (for further information, we refer to the Methods section). (a) Comparison of predicted binding labels versus their experimentally validated results. (b) The left flow plot represents untransduced Jurkat cells, and the right flow plot represents TCR-transduced Jurkat cells showing the expression of the CD3 marker. (c) Flow plot illustrating a true positive TCR binding event, visualized as a double-positive population binding both Tetramer-APC and Tetramer-BV421.

### Functional validation of filtered TCRs

Having shown that our filtering approach removes plausibly noisy data points, we next validated it experimentally. For each of three peptides, we selected ten TCRs: five predicted to be binders (i.e., correctly labeled) and five predicted to be non-binders (i.e., mislabeled). We then functionally assessed the binding specificity of the resulting 30 TCRs. We chose TCRs specific to the peptides TTDPSFLGRY, YVLDHLIVV, and IVTDFSVIK and randomly selected five filtered and non-filtered TCRs. Briefly, TCRs were transduced into TCR KO CD8+ Jurkat cells using TCR-encoding lentivirus and stained for their cognate pMHC antigen (for more information, we refer to the Methods section).

The results of the validation of the 30 TCRs are shown in Figure 6 and Supplementary Figure 7. Overall, the results show a good agreement between the predicted and experimentally derived labels, with 19 out of the 30 (63%) labels being correctly predicted. However, focusing on the predicted noise label, this number increases to 80% (12 out of 15). In contrast, only 47% (7 out of 15) data points predicted to be true binders demonstrated binding. These results demonstrate that while some data points predicted to be true binders cannot be validated, almost all data points that were predicted to be noise are indeed non-binders, showcasing NetTCR’s high specificity in TCR-pMHC binding prediction.

## Discussion

In this study, we introduced a simple, cross-validation-based label noise prediction scheme to identify and filter low-quality data points, and demonstrated its ability to refine public domain training data and improve the biological alignment of NetTCR. While our approach was capable of separating validated from non-validated TCRs, its efficacy differed significantly between data subsets, where the denoising performance was notably poorer for the GLC peptide compared to that of YLQ. This observation is in line with the findings in the original TCRvdb work, where different approaches for separating validated versus non-validated TCRs consistently displayed a higher performance for YLQ compared to GLC. As mentioned in our results, the bimodal separation of scores for YLQ may be driven by high sequence similarity among its validated TCRs, creating a strong, easily identifiable signal.

A notable limitation of our proposed denoising approach is its inability to separate epistemic from aleatoric uncertainty. Aleatoric uncertainty represents true label noise; data points that are fundamentally incorrect due to experimental artefacts. Epistemic uncertainty, however, arises from the model’s own uncertainty in regions of the TCR-pMHC sequence space where training data is sparse or contains convoluted signals, which is the case for data for many peptides. By filtering positive examples that score below a stringent threshold derived from the negative data score distribution, we risk discarding genuine but hard-to-learn TCR-pMHC pairs alongside truly mislabeled data. While we attempt to address this issue through ensemble voting, which may reduce the epistemic component, cases where insufficient training data is available to accurately learn interaction motifs certainly remain [33].

Using structural modeling confidence for models generated with AlphaFold 3 serves as an orthogonal confirmation of our findings and shows that data predicted to be noise by our denoising scheme, generally, is modeled with low confidence. Interestingly, we see a significant correlation between the proportion of data denoised for a given peptide and the average modeling confidence of AlphaFold, indicating a clear concordance in how the two approaches score the same data (Supplementary Figure 4). However, for the most overrepresented peptides, RVR, AVF, and KTF, the confidence score delta between filtered and non-filtered is small, despite AlphaFold scoring some complexes in the 0.8 to 0.9 confidence range. This suggests that NetTCR in these cases is unable to accurately detect the biological signal in the data. This is likely because the signal-to-noise ratio for these peptides is too low to train a robust NetTCR model despite ample training data, which consequently makes the model incapable of confidently separating clean and noisy samples (Supplementary Figure 1) [5]. This suspicion is further corroborated by the increasing delta between filtered versus non-filtered confidence as peptides become less represented in filtered data, suggesting that for peptides of higher data quality, NetTCR is more robust to noise (Figure 4b).

The experimental validation of the denoising approach on 30 selected TCRs demonstrated that predictions of noise were highly reliable, with 80% of the TCRs classified as noise being confirmed as non-binding. This finding suggests that the proposed framework achieves high specificity for identifying likely misannotated TCR-pMHC interactions. In contrast, 47% of TCRs predicted to be true binders were experimentally confirmed. This may in part be explained by low-affinity TCR-pMHC interactions not being readily identified by pMHC tetramer staining and fluorescence. Further, spurious signals existing in the noisy training data subsets for these peptides could bias NetTCR toward scoring noise-like complexes as binders. Consequently, experimental validation on larger data sets is needed in order to provide more solid evidence for the power of the suggested denoising approach.

The limited benchmark dataset for evaluating the denoising performance of our approach highlights the current challenge for the TCR specificity prediction task: there is a significant lack of gold standard data [3], [34]. This inhibits model development, as we are evaluating predictive performance on data deviating from the underlying biological rules governing TCR-pMHC interaction. Particularly, we highlight this when demonstrating the increased performance delta of NetTCR for validated versus non-validated positive data points. This lack of reliable data also creates challenges in developing data denoising methods, as basing model development on small subsets of data often leads to poor generalization, particularly when considering TCR, antigen, and interaction mode diversity [35], [36]. Until more reliable and diverse data become available, orthogonal methods not reliant on supervised training on TCR specificity data, such as AlphaFold-based approaches, appear as good alternatives despite their shortcomings.

Given these limitations, our findings nevertheless demonstrate that supervised learning models can be used not only for TCR specificity prediction, but also as effective tools for identifying potentially mislabeled TCR-pMHC interactions. As larger functionally validated benchmark datasets become available, such approaches may play an important role in improving both the quality of training data and the reliability of computational TCR specificity prediction.

## Data availability statement

The datasets and code used in this research can be accessed at our GitHub repository https://github.com/mnielLab/NetTCR-torch/.

## Acknowledgements

Research reported in this publication was supported by the Novo Nordisk Foundation Data Science Collaborative Research program under the grant NNF24OC0089619 and by the Innovative Medicines Initiative 2 Joint Undertaking under grant No 101007799 (Inno4Vac).

## Notes

### Competing Interest Statement

The authors have declared no competing interest.

