## Supplementary for "Exploring noise in TCR specificity data through supervised learning"

#### Number of positive examples in the NetTCR training dataset and predictive performance

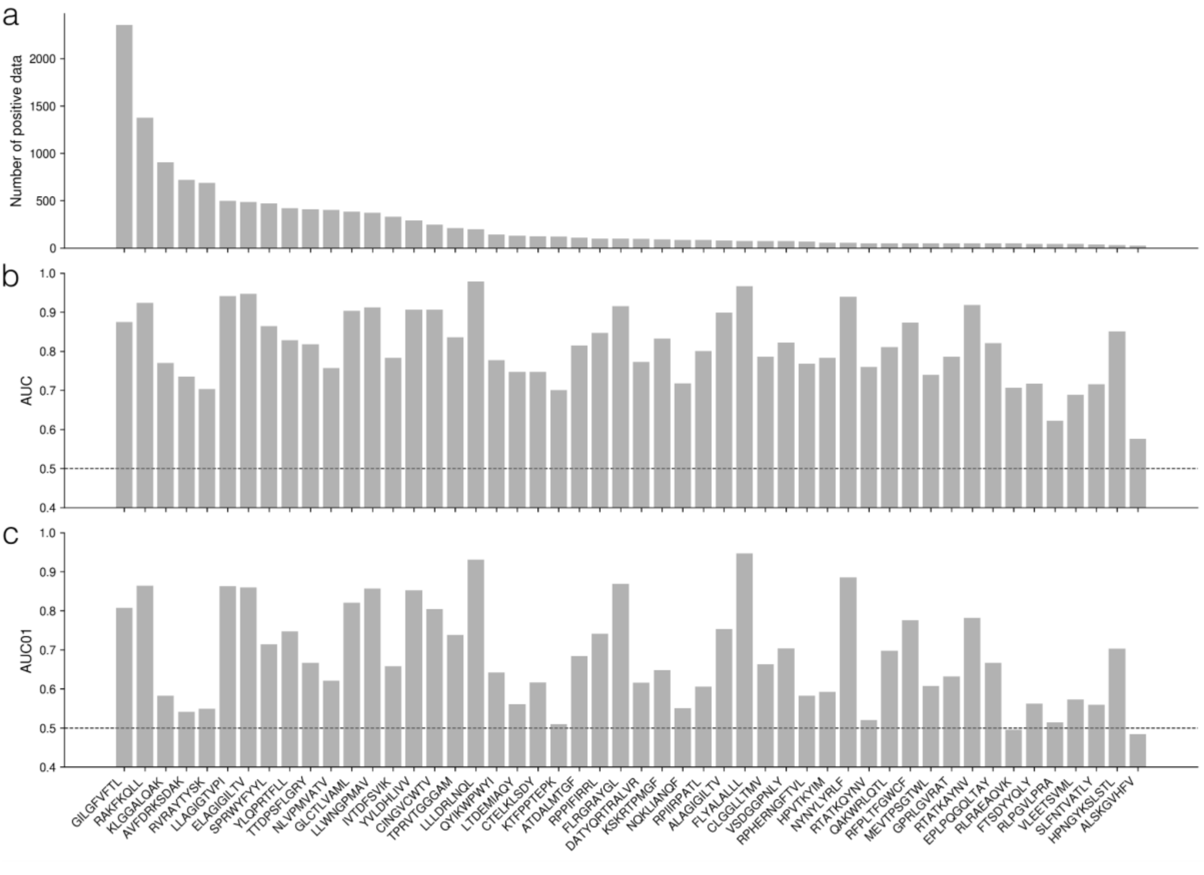


**Supplementary Figure 1: The number of positive data points and predictive performance for each peptide in the NetTCR training dataset.** (a) Number of positive data points for each peptide. (b) and (c) Predictive performance per peptide. For more information on how the dataset was constructed and how NetTCR was used to score the data, we refer to the methods section.

#### YLQ and GLC TCR sequence similarity

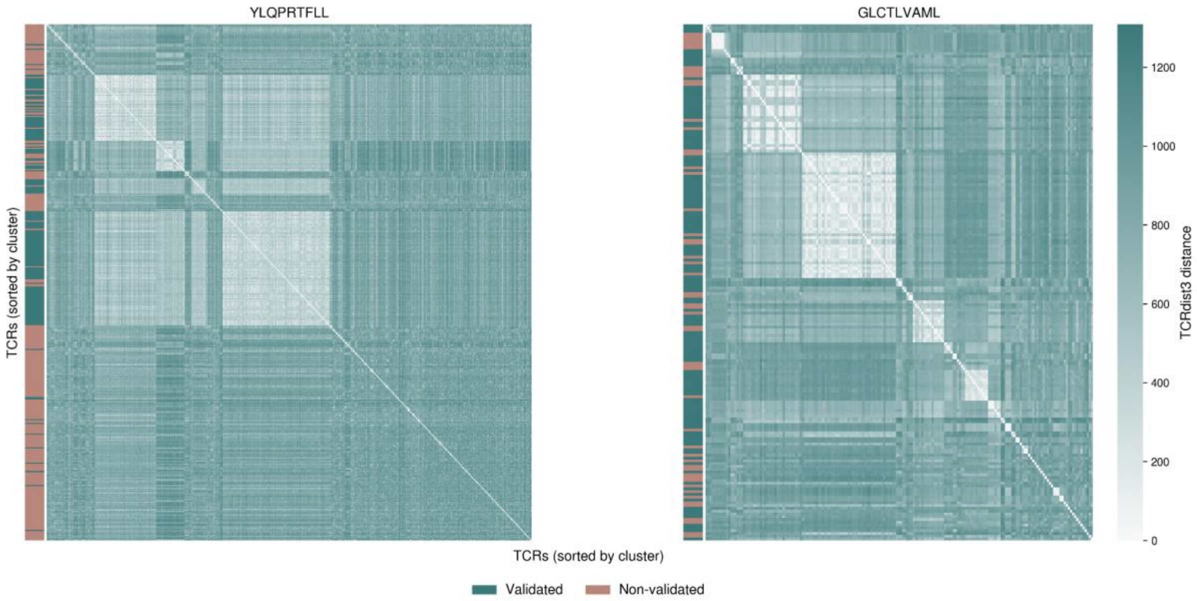


**Supplementary Figure 2: TCRvdb TCR sequence similarity.** Sequence distances for YLQ and GLC TCRs from the TCRvdb were computed with TCRdist3 (1). Weights for CDR1, CDR2, and CDR2.5 were set to 1, and the weight for CDR3 was set to 3. For a given TCR, α- and β-chain distances were summed to get a single distance metric per TCR. The distance matrices were clustered using the AgglomerativeClustering method from scikit-learn, using average linkage and a distance threshold of 500. The clustering was used to sort the distance matrix visualization (2).

### Denoising performance metrics for YLQ and GLC

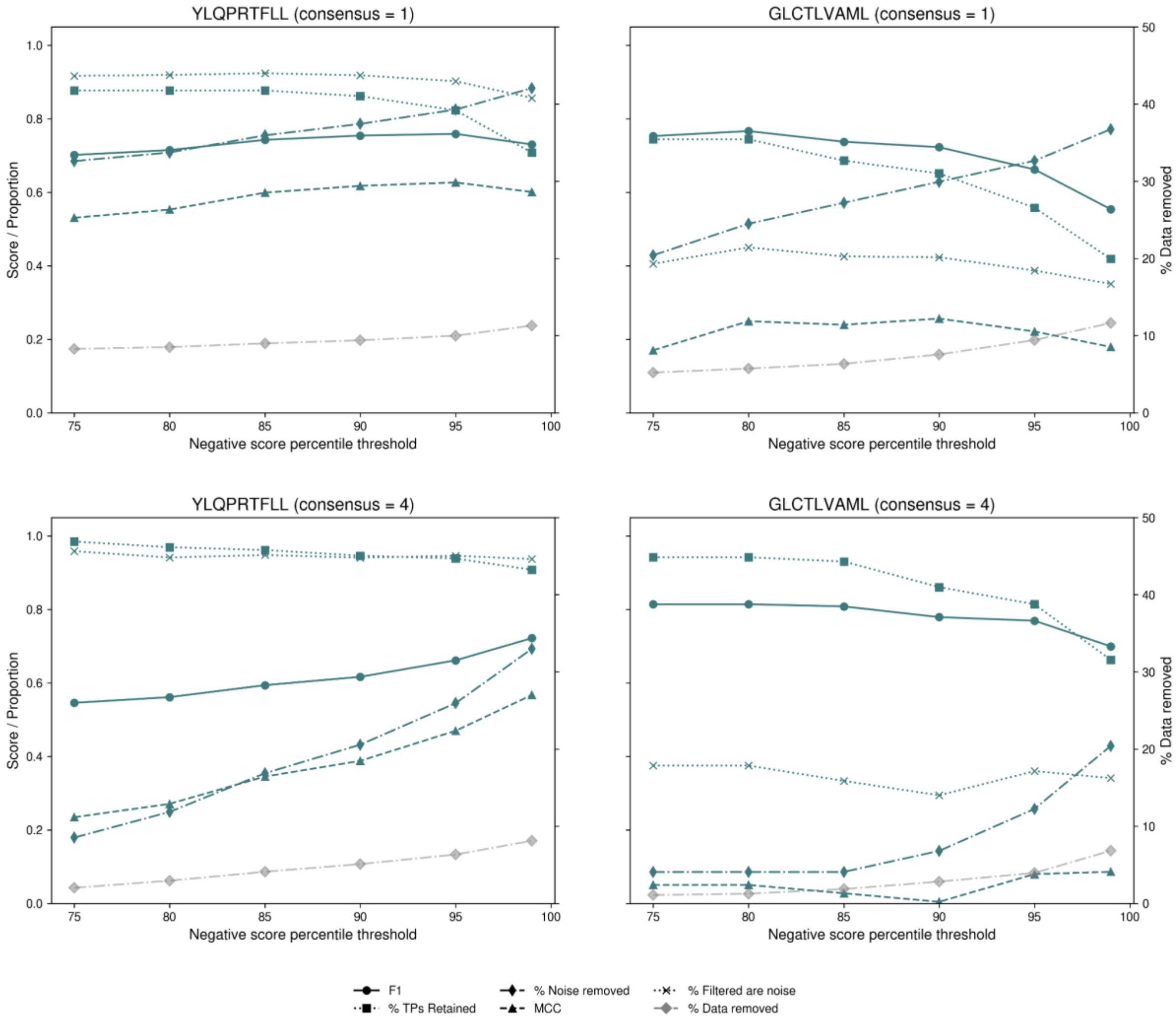


**Supplementary Figure 3: Congruence of filtered examples and their validation status in the TCRvdb dataset for different consensus counts.** A NetTCR mode was used to score positive TCRs towards the two peptide examples in the NetTCR dataset. For each peptide, the score threshold corresponding to the Nth percentile of negative TCR examples was calculated, and all positive examples with a predicted score less than this threshold were labeled as considered to have a noisy label and were therefore removed from the dataset. A higher score threshold consequently results in more stringent filtering. Based on the noise label, a confusion matrix was constructed with the number of true positives (validated TCRs not labeled as noise), false negatives (validated TCRs labeled as noise), true negatives (non-validated TCRs labeled as noise), and false positives (non-validated TCRs labeled as not noise). Thresholds were computed for the 75th, 80th, 85th, 90th, 95th, and 99th percentiles of negative data point scores, and the MCC and F1 for classifying validated versus non-validated were computed for each peptide from the confusion matrix generated at each threshold for datasets filtered using each threshold. Data points were only considered to be noise if all nested cross-validation models agreed in their predictions.

### Correlation between AlphaFold 3 confidence and filtering enrichment

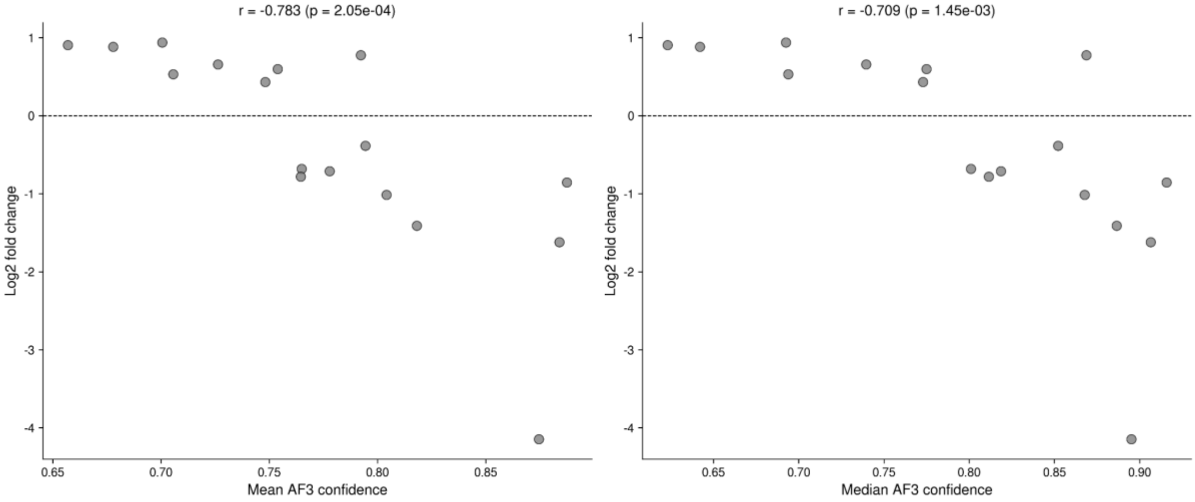


**Supplementary Figure 4: Mean and median per-peptide AlphaFold 3 confidence are correlated with Log2 fold change of filtered peptides.** TCR-pMHC complexes for peptides significantly enriched or depleted in filtered data were modeled with AlphaFold 3. Scores were max-pooled per complex, and the median and mean confidence values per peptide were computed. Computing the Pearson correlation between these values and the Log2 fold change of peptide frequencies in filtered versus all data reveals a significant relationship between predicted noise proportions and AlphaFold confidence values ($p$=2.02e-04 and $p$=1.45e-03).

AlphaFold 3 based scoring of YLQ and GLC

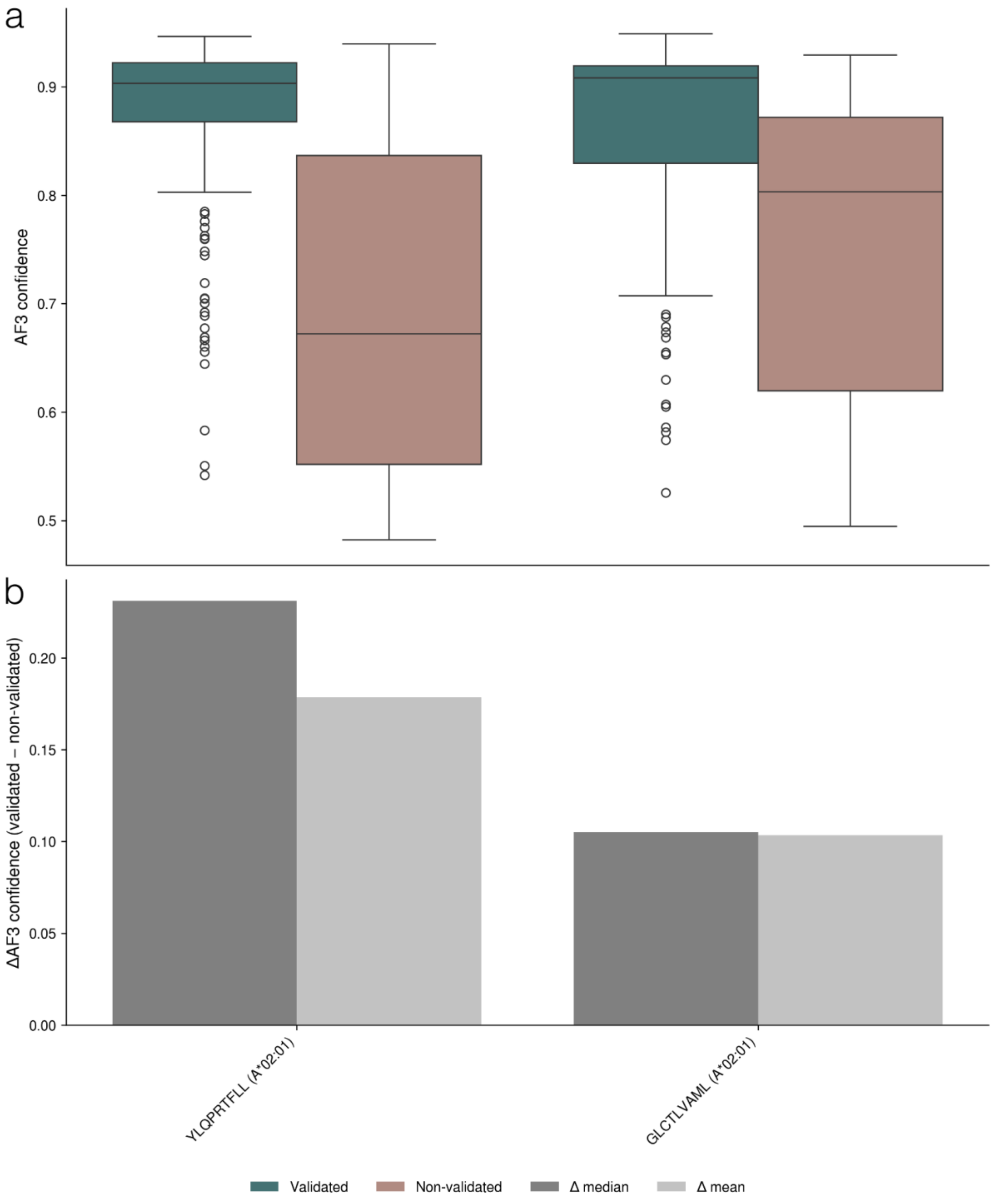


**Supplementary Figure 5: AlphaFold 3 confidence scores of validated and non-validated TCRs.** TCR-pMHC complexes for peptides YLQ and GLC were modeled and scored using the AlphaFold confidence metric (for more information, we refer to the methods section). (a) Distribution of confidence scores for validated and non-validated examples. (b) The difference in mean and median confidence values of validated and non-validated examples. These results show that AlphaFold accurately separates validated and non-validated TCRs, corroborating its use as a secondary benchmark for our denoising approach.

#### Predictive performance of denoised models on TCRvdb data

We examined how filtering data affects the behaviour of NetTCR by denoising the NetTCR training dataset and retraining NetTCR for all score thresholds. Note that in this case, no filter voting system was used. As expected, models trained on denoised data exhibit higher confidence in their predictions as ambiguous examples are removed from the training data. However, quantifying this change in terms of ROC-AUC for predicting validation status shows only a minor increase in AUC and AUC01 for GLC, and a decrease in AUC01 for YLQ. This observation reflects the fact that the baseline model, to a very high degree, is robust to noise and, in particular, for YLQ, already separates the classes well.

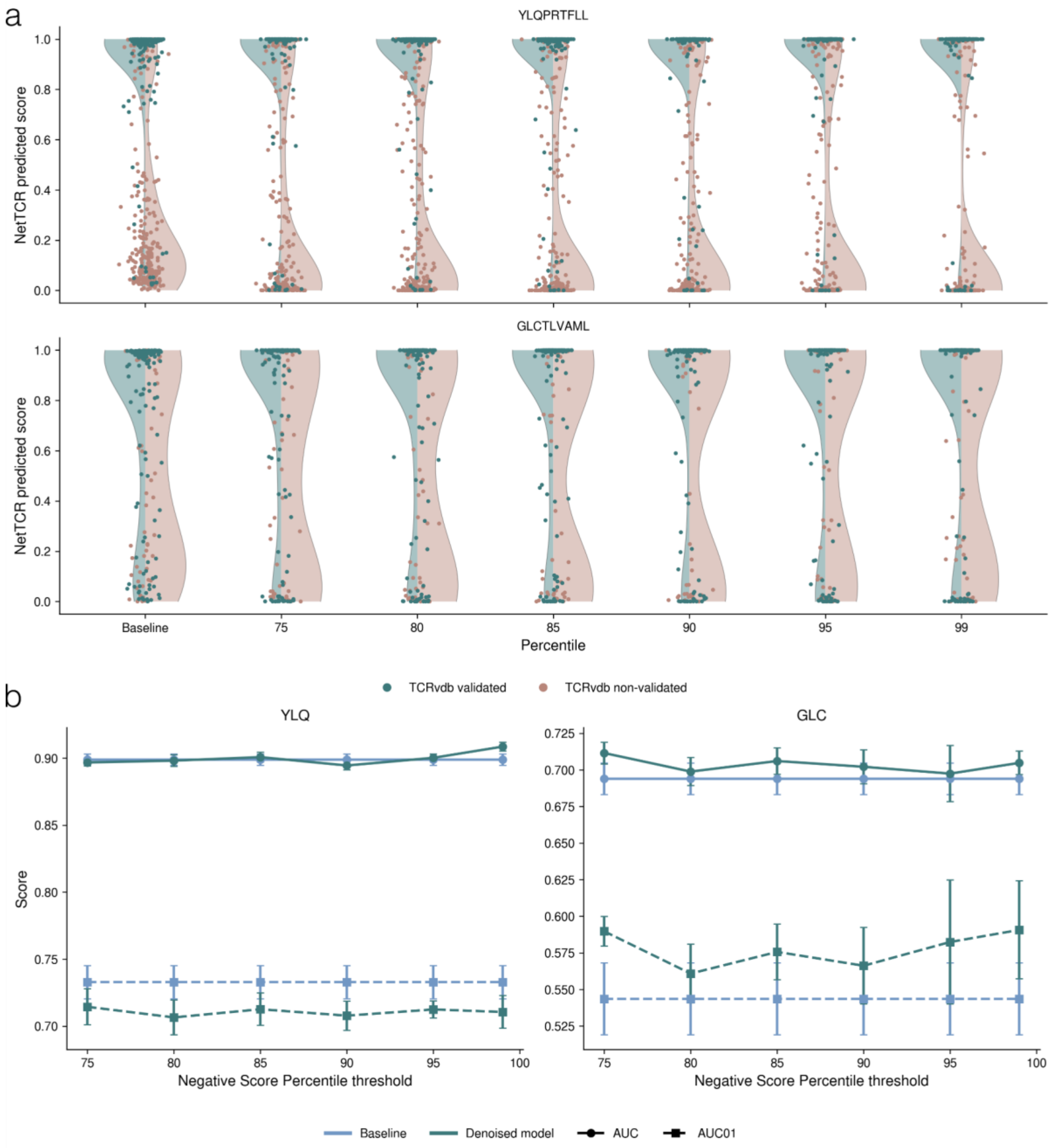


**Supplementary Figure 6: Performance for predicting validated and non-validated status for data points in the TCRvdb dataset.** NetTCR models were trained on NetTCR datasets denoised using score thresholds corresponding to either the 75th, 80th, 85th, 90th, 95th, or 99th percentile of predicted scores for negative data points. The NetTCR models trained on the denoised datasets were then used to assign prediction scores for the TCRvdb datasets using cross-validation for each score threshold. (a) NetTCR score distributions for each score threshold used to filter training data. (b) AUC and AUC01 for predicting validation status versus the score threshold used to filter training data. Models were trained using ten different parameter initializations.

### Results for experimental validation of predicted noise

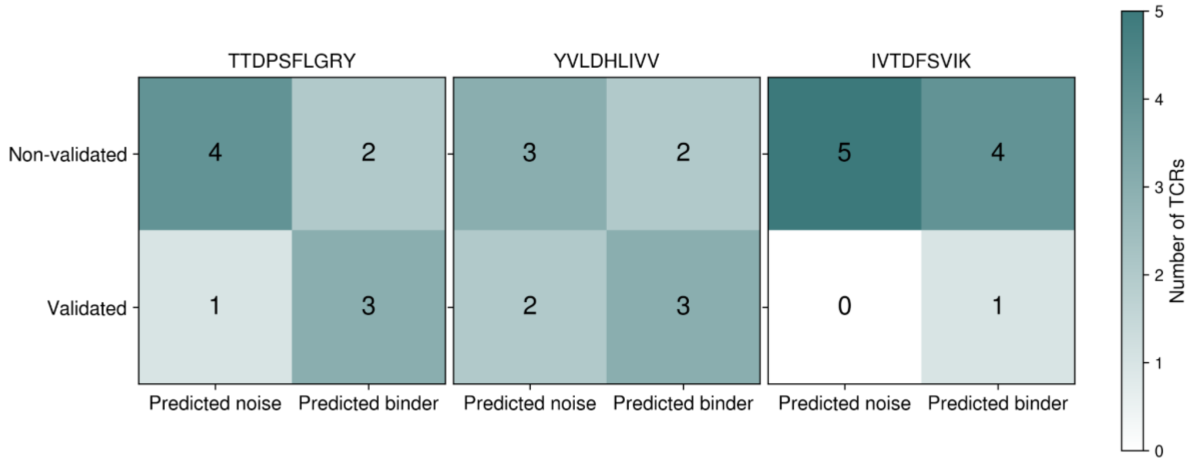


**Supplementary Figure 7: Experimental validation of TCRs predicted to be noise.** 30 TCRs from public databases for peptides TTDPSFLGRY, YVLDHLIVV, and IVTDFSVIK were scored by NetTCR to be either binders or non-binders. TCRs were transduced into TCR KO CD8+ Jurkat cells using TCR-encoding lentivirus and stained for their cognate pMHC antigen, and their binding status was compared to their predicted labels (for further information, we refer to the Methods section).
